# Super-resolution mass spectrometry enables rapid, accurate, and highly-multiplexed proteomics at the MS2-level

**DOI:** 10.1101/2022.07.29.501912

**Authors:** Anton N. Kozhinov, Alex Johnson, Konstantin O. Nagornov, Michael Stadlmeier, Warham Lance Martin, Loïc Dayon, John Corthésy, Martin Wühr, Yury O. Tsybin

## Abstract

In tandem mass spectrometry (MS2)-based multiplexed quantitative proteomics, the complement reporter ion approaches (TMTc and TMTproC) were developed to eliminate the ratio-compression problem of conventional MS2 level approaches. Resolving all high *m/z* complement reporter ions (∼6.32 mDa spaced) requires mass resolution and scan speeds above the performance levels of Orbitrap™ instruments. Therefore, complement reporter ion quantification with TMT™/TMTpro™ reagents is currently limited to 5 out of 11 (TMT) or 9 out of 18 (TMTpro) channels (∼1 Da spaced). We first demonstrate that a Fusion™ Lumos™ Orbitrap™ can resolve 6.32 mDa spaced complement reporter ions with standard acquisition modes extended with 3-second transients. We then implemented a super-resolution mass spectrometry approach using the least-squares fitting (LSF) method for processing Orbitrap transients to achieve shotgun proteomics-compatible scan rates. The LSF performance resolves the 6.32 mDa doublets for all TMTproC channels in the standard mass range with transients as short as ∼108 ms (Orbitrap resolution setting of 50 000 at *m/z* 200). However, we observe a slight decrease in measurement precision compared to 1 Da spacing with the 108 ms transients. With 256 ms transients (resolution of 120 000 at *m/z* 200), coefficients of variation are essentially indistinguishable from 1 Da samples. We thus demonstrate the feasibility of highly-multiplexed, accurate, and precise shotgun-proteomics at the MS2 level.

## Introduction

Multiplexed quantitative proteomics based on tandem mass spectrometry (MS2) is most commonly realized using peptides covalently labeled with isobaric tandem mass tags (iTRAQ, TMT, TMTpro).^1-4^ It may simultaneously probe up to 18 sample-specific biological conditions.^5^ The analytical benefits include the inherently high measurement precision, omission of missing values, and reduced experimental time per proteome. The corresponding applications have been exploited for basic and applied biological research, including translation regulation,^6^ embryology,^7^ breast cancer treatment,^8^ and lung cancer metastasis.^9^

The TMT isobaric reagents consist of a reactive moiety that covalently binds to a peptide and reporter and normalization parts that uniquely encode several quantitation channels with different numbers of heavy isotopes – ^13^C and ^15^N.^2,3,10^ The reporter parts are detected as singly charged reporter ions in the low, *m/z* 110 – 140, mass range. The abundances of the reporter ions quantify peptides and associated proteins in each proteome. The reporter ion cluster contains channels separated by ∼1 Da from other channels and doublets containing two channels encoded by the ^13^C^14^N vs. ^12^C^15^N isotopes, which differ by only ∼6.32 mDa.^11^ To baseline resolve the 6.32 mDa doublets in this (low *m/z*) mass range, a resolution value of about 60 000 is sufficient. This level of resolution performance is readily provided by modern Orbitrap™ Fourier transform mass spectrometers (FTMS) at their resolution setting of 50 000 at *m/z* 200.

Despite its attractiveness, the MS2-level multiplexed proteomics suffers from measurement artifacts due to co-isolation and co-fragmentation of co-eluting peptides when applied to complex samples. The MS3-level tandem MS approaches have been developed to address this limitation.^12-14^ Adding one more MS step mitigates the peptide co-isolation issue but reduces sensitivity and throughput.^15^

An alternative approach to tackle peptide co-isolation and the associated reporter ion ratio-compression problem is MS2-level complement reporter ion quantification.^16-19^ Following this method, the species to be monitored are the complementary reporter ions – the remainders of the precursor ions labeled with the normalization parts and the reactive groups. For TMTpro/TMT reagents, the complementary ion clusters will have one charge less than the isolated precursor ions and thus will be detected in an even higher mass range, up to 2 000 *m/z*, and sometimes beyond. The Orbitrap mass analyzer baseline resolves the complementary ion channels that differ by ∼1 Da at a proteomics-grade scan rate. However, this reduces the number of encodable channels from 11 to 5 when using TMT and from 18 to 9 when using TMTpro.^18,19^

Adding complementary reporting channels (TMTc/TMTproC clusters) with the 6.32 mDa doublets to the analysis increases the number of quantification channels but requires resolution performance that is not attainable even with the state-of-the-art Orbitraps. For example, to baseline resolve the 6.32 mDa doublets of the complementary reporter ions at *m/z* 2 000, a resolution value of 600 000 is necessary (which corresponds to a virtual resolution setting of 2 000 000 at *m/z* 200). However, this capability is not available for general Orbitrap users.^20,21^ On the other hand, user access to extended-duration transients can be readily provided using external high-performance data acquisition (DAQ) systems.^22-24^ However, even if available, such long transients would drastically reduce the MS scan rates and are thus prohibitive for routine shotgun high-throughput proteomics applications using liquid chromatography (LC)-MS analysis. In addition, extended periods of ion oscillation in the mass analyzer may result in the loss of ion motion coherence, manifesting itself through pronounced transient decay and frequency shifts.^21,25,26^ Finally, the FT-type signal processing methods, being the *spectral* estimators (as they yield profile-type data with the characteristic peak shape), are prone to frequency (*m/z*) and amplitude (ion abundance) errors of peak maxima.^27^

The super-resolution (SR) algorithms for transient data processing were introduced for FTMS to overcome the abovementioned limitations.^28^ Compared to FT-based approaches, a SR method is characterized by a less strict uncertainty principle for the resolution performance as a function of the detection period and may provide the same resolution as the FT methods using multiple-fold shorter transients.^27^ Additionally, the difference in the uncertainty principles of the SR vs. FT methods reduces the peak *m/z* and amplitude measurement errors; this may translate to improved accuracy of quantitation, particularly in the TMTc/TMTproC doublets analysis. Previously, we implemented the least-squares fitting (LSF) of time-domain signal processing as a powerful SR method for FTMS.^29^ By definition, the LSF method is a targeted method that requires information on the (estimated) frequency of interest (*m/z* values) as an *a priori* knowledge. The latter matches the case of multiplexed quantitative proteomics for both low *m/z* reporter and complementary ion detection. The output of LSF processing contains the amplitudes and refined frequencies of ion signal components in the transient (and, thus, the abundances and *m/z* values of peaks in the peaklist-type mass spectrum). Here, we combined the LSF capabilities to resolve the nearly isobaric ion signals using shorter time-domain transients with the highly-plexed TMTc/TMTproC workflow’s ability to overcome the inherent MS2-level TMT/TMTpro strategy limitations.

## Methods

### Sample preparation

Yeast peptides from *Saccharomyces cerevisiae* cell lysates were prepared as described previously^18,30^ and labeled with selected TMTpro reagents to yield 12 complement reporter ion channels (four 6.32 mDa doublets and four 1 Da singlets) or with selected TMT reagents to yield four channels (one 6.32 mDa doublet and two 1 Da singlets). In the case of TMTproC labeling, only 12 out of 18 channels were employed due to the degeneration (overlap) of the complement reported ions. Using only four channels for the TMTc approach validation was sufficient to evaluate method performance and applicability.

### Mass spectrometry

LC-MS experiments were conducted on two similar Fusion™ Lumos™ Orbitrap™ FTMS instruments (Thermo Fisher Scientific, San Jose, CA, USA). Time-domain signals of various standard and extended durations, up to 3 seconds, were acquired using an external high-performance DAQ system (FTMS Booster X2, Spectroswiss, Lausanne, Switzerland) in parallel to RAW file acquisition. The FTMS Booster X2 is an add-on to diverse Orbitrap models, with the previously reported implementation on the Q Exactive Orbitraps.^22-24^

Notably, the employed DAQ system enables the acquisition of *phased* transients, i.e., transient signals whose sinusoidal components induced by the individual *m/z* ion rings trapped in the mass analyzer are all nearly in-phase. In order to receive the analog transient signals from the Orbitrap pre-amplifier’s output, the external DAQ system was interfaced through its two high-impedance analog inputs (i.e. a single differential input) to the two outputs of the Orbitrap pre-amplifier using T-splitters (Figure S1, Supporting Information), as described elsewhere.^23,24^

To increase resolution beyond conventional FTMS, the acquisition of longer transients was enabled by introducing “dummy” scans into the experimental logic without hardware or operational software modifications (Figure S2, Supporting Information).^23,24^ As a result, long time-domain transients, up to 6 s in this work, could be acquired.

### Data processing

Assignment of MS^2^ spectra was performed using the SEQUEST algorithm^31^ by searching the data against the Uniprot *Saccharomyces cerevisiae* proteome. A peptide-level MS^2^ spectral assignment false discovery rate of 1% was obtained by applying the target decoy strategy with linear discriminant analysis as described previously.^32^ Super-resolution processing, absorption-mode FT, and calculations of reference *m/z* values for TMTc and TMTproC ions were performed using the Peak-by-Peak software package (Spectroswiss) running on one or several 8-core desktop computers, eachequipped with 32 GB RAM and a compute unified device architecture (CUDA)-compatible graphics card (GPU) of the compute capability index ranging from 5.2 to 6.1. The super-resolution data processing was performed using the LSF approach similarly to the previously described fundamentals and implementation.^29^ The basis functions of the LSF method correspond to the TMTproC (or TMTc) cluster and compose of the individual channels (∼1 Da spacing) and doublets (∼6.32 mDa spacing), Figure S3, Supporting Information. In addition to the general accessibility of the LSF-like methods in the contemporary data processing tools, including Python and MATLAB, the LSF calculations module for TMT-based applications is now a part of the Peak-by-Peak software. For the use-case with multiple computers in this work, individual instances of Peak-by-Peak launched on each computer form a cluster network for parallel processing of an LC-MS run.

The computational overhead imposed by the LSF processing significantly exceeds the FT times, as expected. Nevertheless, owing to the parallelizable natures of the LSF task (parallel execution for a single transient on a GPU) and the LC-MS run (parallel execution for multiple transients, e.g., on N computers each with a GPU), the total LSF processing time per LC-MS run can be adjusted to be comparable to (or less than) the LC-MS run time. For instance, with the data processing set-up of this work, for the 2-hour LC-MS runs with MS2 scans acquired at resolution settings of 50 000 and 120 000, the LSF processing times were measured to be 2.9 hr/N and 7.2 hr/N, respectively, where N is the number of employed computers (GPUs), e.g. 1-4. Overall, acceleration of LSF processing of LC-MS runs is possible by increasing the number of GPUs and employing GPUs of greater performance (a larger CUDA compute capability index).

The outcome of the Peak-by-Peak data processing was extracted in the form of the mzXML files and uploaded to the GFY software licensed from Harvard for further analysis^33^. Peptides were only considered quantified if the average signal to the equivalent FT noise ratio (S:N) of the complementary ions was greater than 20 (240 total S:N). Pipetting errors were corrected by dividing the signal in each complementary ion channel by the channel’s median S:N ratio across all peptides. For the singlets, peptide coefficients of variations (CVs) were calculated by taking the population standard deviation of the S:N ratios of the four channels for each peptide and dividing by the mean complementary ion S:N. Doublet CVs were calculated identically on four randomly selected channels for each peptide.

## Results

### TMTc/TMTproC method validation for Orbitrap FTMS

Practical usage of the highly-plexed TMTc and TMTproC workflows with Orbitraps requires achieving UHR performance in a wide mass range. According to the Orbitrap fundamentals, resolution in a mass spectrum for a given transient length reduces as a square root of *m/z*. In addition, for the 6.32 mDa doublets in TMTc/TMTproC clusters, higher charge states at a given value of *m*/z impose stricter resolution requirements. **Figure 1** estimates the theoretical Orbitrap performance to baseline resolve a 6.32 mDa doublet in a TMTc/TMTproC cluster as a function of the doublet’s *m/z* and the precursor’s charge state. The singly charged doublet (for a doubly charged precursor) detected at *m/z* ∼1900 requires at least a 3.7 s transient to be baseline resolved. The same length transients will baseline resolve the doubly charged doublet (for a triple charged precursor) only up to about *m/z* 1300.

**Figure 1.**
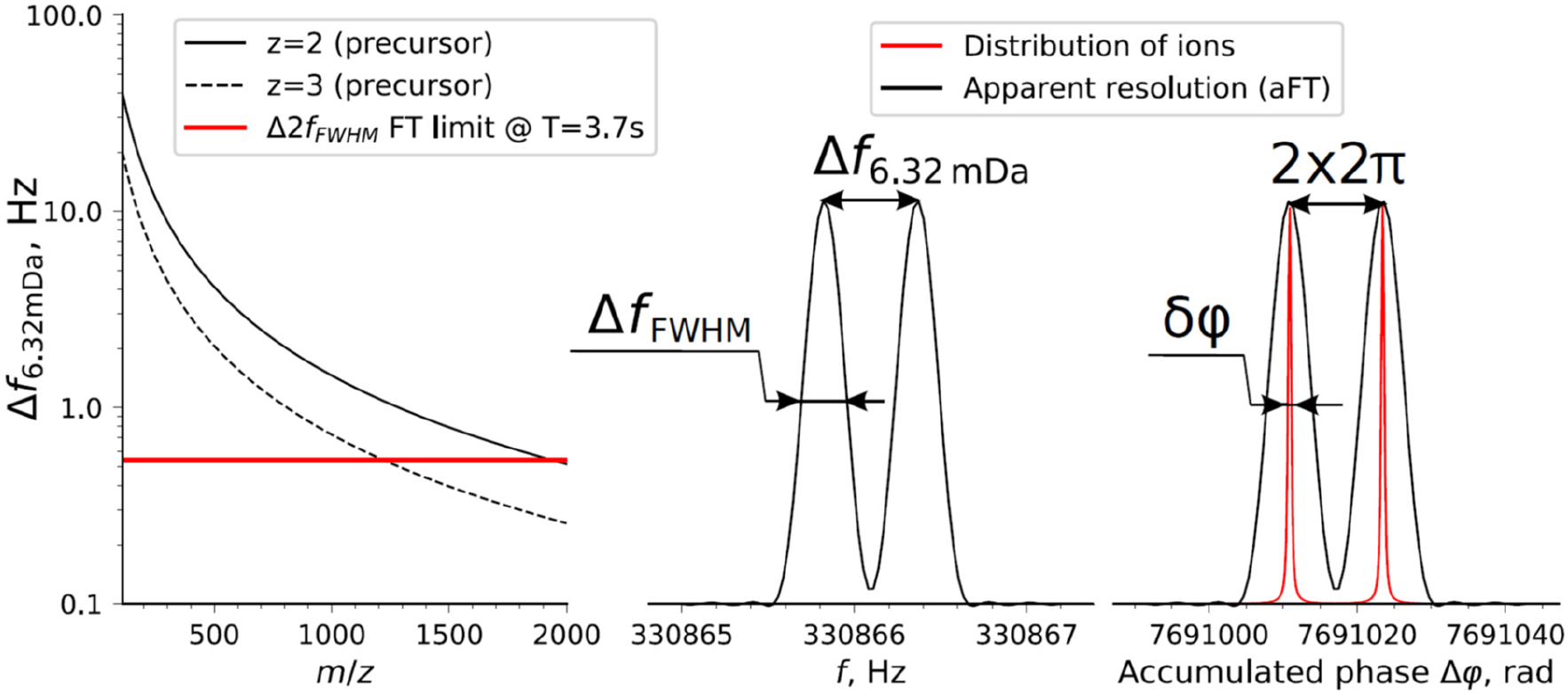
The uncertainty principle of Fourier transform (FT), represented for the spectral components corresponding to 6.32 mDa doublets in complement reporter ion clusters. Left panel: the frequency difference for a doublet as a function of the doublet’s *m/z*. The FT resolution (soft) limit, T = 3.7 s, is shown by example (it corresponds to the doublet at *m/z* ∼1900, 2+ precursor). Center panel: apparent FT resolution for the doublet in question, at T = 3.7 s. Right panel: comparison of the apparent FT resolution (in the phase scale) shown in black and the fundamental limit to ion separation (distribution of ions over their total phase accumulated during ion detection) shown in red.

The analysis of the UHR mass spectra is thus important to reveal the structure of the TMTc/TMTproC clusters for high mass range and validate the method’s applicability. The latter involves considerations for the space-charge-induced coalescence between the quantitation channels in the doublets. However, the capability to acquire the UHR mass spectra produced by FT of 3.7 s time-domain signals cannot be accessed by a general Orbitrap user. On the other hand, recent innovations in high-performance data acquisition electronics and allied digital signal processing have enabled the acquisition and processing of extended-period time-domain signals from Orbitrap instruments. It should be noted that the experimental settings and manufacturing quality of a given Orbitrap platform may influence the parameters of the acquired time-domain signals and thus the practical ability to reach the UHR performance with accurate estimates of ion signal intensities and frequencies. The critical quality attributes of time-domain signals include the linearity of their phase function as well as signal decoherence effects (including transient decay and frequency shifts) for long ion detection events.

We first enabled the UHR Orbitrap FTMS operation through an external high-performance DAQ system (FTMS Booster X2). This set-up was employed to analyze isotopically-labeled samples with 4 TMTc channels with a single 6.32 mDa doublet, **Figure 2**. Acquisition and processing of the 3 s transients using absorption-mode FT (aFT) showed an unmodified ultra-high-field D20 Orbitrap mass analyzer (as employed in Orbitrap™ Fusion™ Lumos™) could resolve the 6.32 mDa-spaced TMTc peaks in the required *m/z*-range up to 2000 *m/z* (Figure S4 and Figure S5, Supporting Information). Expectedly, the maximum manufacturer-supported resolution of this particular Orbitrap model provides a close to the baseline resolution of the singly charged TMTc clusters up to *m/z* 800, but cannot baseline resolve them in the higher *m/z* range (Figure S4 and Figure S5, Supporting Information). These results confirm the fundamental experimental feasibility of Orbitrap FTMS to perform the highly-plexed TMTc approach in a wide mass range. Nevertheless, in addition to the low throughput of UHR Orbitrap experiments, the measured relative abundances exhibited relatively high CVs for both 6.32 mDa and 1 Da spacing, demonstrating that some detrimental effects develop over the very long ion detection times, Figure S6, Supporting Information.

**Figure 2.**
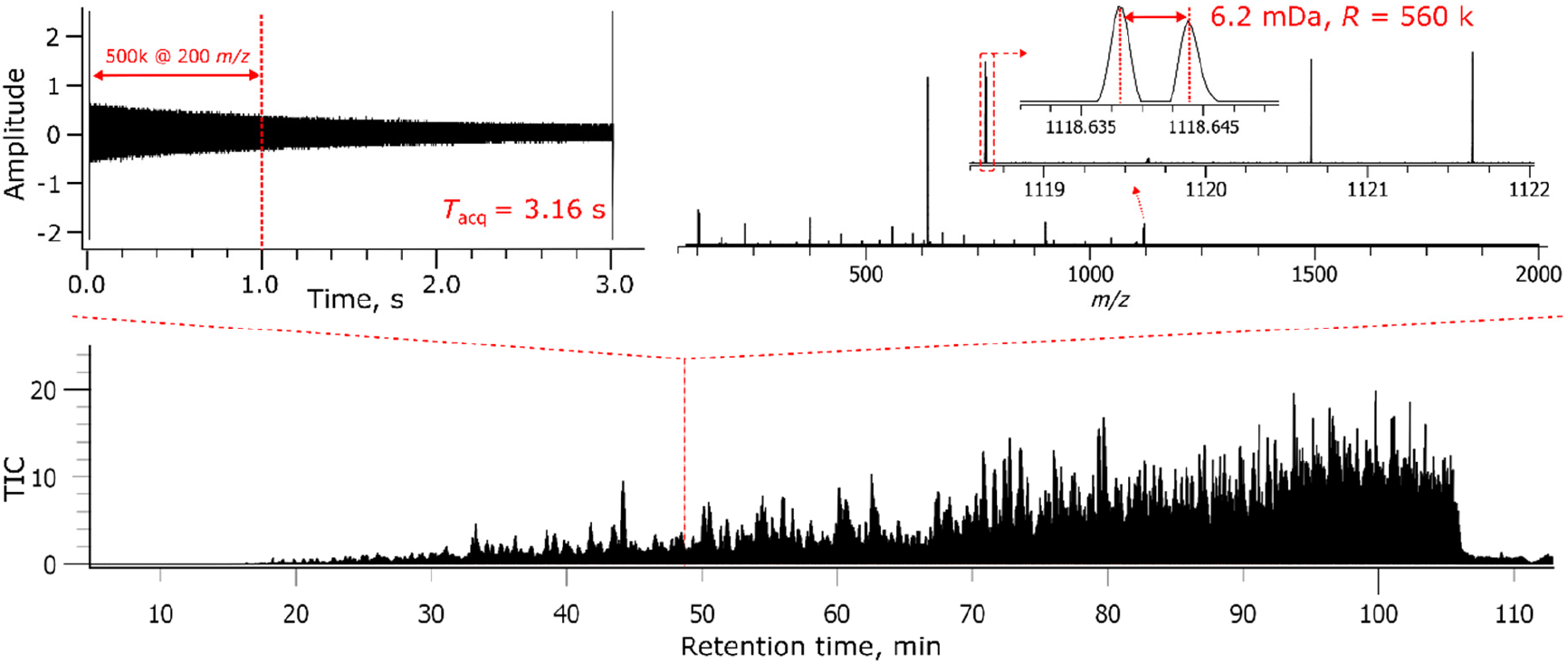
Illustration of aFT signal processing with extended detection period data acquisition in the LC-MS analysis of a 4-plex TMT labeled sample with equal concentrations over the 4 TMTc channels. The LC-MS experiment was performed with an Orbitrap Fusion Lumos FTMS instrument, with MS2 data acquired at the resolution setting of 500 000 at *m/z* 200. The corresponding >3 seconds long transients were acquired in parallel using an external high-performance data acquisition system (FTMS Booster X2) and a user-defined Orbitrap experimental sequence method (Figure S3). This length is equivalent to a virtual resolution setting of 1.5 million at *m/z* 200.

Therefore, enabling highly-plexed complement ion quantification requires significantly shorter time-domain transients. As a proof of concept, the original LSF processing was applied to the shortened time-domain transients for the same 4-plex TMTc model sample. This initial example demonstrates that the 6.32 mDa doublets could be resolved with a transient period reduced to 512 ms (resolution setting of 240 000 at 200 *m/z*) using the LSF method, Figure S7, Supporting Information.

### Super-resolution analysis for 12-plex TMTc/TMTproC workflows

The promising initial results of the LSF method application to the 4-plex TMTc workflow have encouraged us to apply the super-resolution analysis to the highly-plexed TMTproC workflow. Several LC-MS experiments were performed on another Fusion Lumos instrument, with MS2 scans acquired at the resolution settings 120 000 (256 ms detection period), 60 000 (128 ms), and 50 000 (∼108 ms). The application of the LSF processing in the analysis of isotopically-labeled samples with 12 channels with four 6.32 mDa doublets confirmed the feasibility of the approach, **Figure 3**. Mass spectral representation in either eFT or aFT for the same dataset does not resolve the doublets. Compared to Figure 2, the LSF parameter optimization allowed the required minimum time-domain transient periods to be shortened by 10-fold to 256 ms (Figure 3). The latter period corresponds to the resolution setting of 120 000 at *m/z* 200, which specifically serves the needs of the highly-plexed quantitative proteomics. This result was achieved provided that certain experimental conditions are met, such as ion-ion interactions in the doublets are sufficiently below the coalescence threshold.

**Figure 3.**
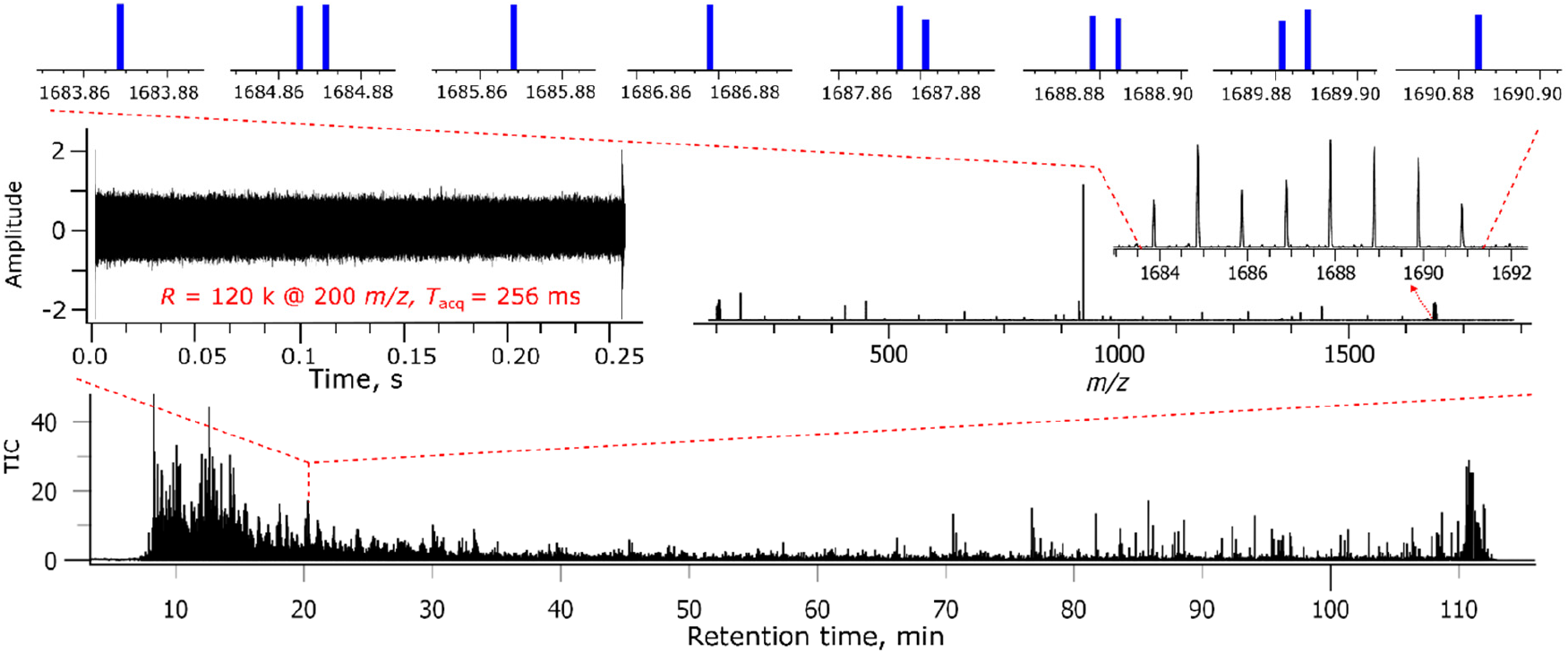
Absorption mode FT (aFT) and LSF analysis of a 12-plex TMTpro™ labeled yeast sample with equal concentrations over the 12 TMTproC channels. The LC-MS experiment was performed with an Orbitrap Fusion Lumos FTMS instrument, with MS2 data acquired at the resolution setting of 120 000 at *m/z* 200. The corresponding 256 ms transients were acquired in parallel using an external high-performance data acquisition system (FTMS Booster X2).

### Quantitation precision of LSF-based 12-plex TMTproC workflows

Resolving the doublets in the whole mass range of TMTc (TMTproC) ions at proteomics-grade scan rates is an important proof-of-concept achievement of the LSF processing, but the obtained quantitation precision needs to be evaluated in detail. The density distributions for the CV values calculated for the doublets and singlets obtained in the LSF analyses of the 12-plex TMTproC experiments with MS2 scans acquired at different resolution settings are shown in **Figure 4**. Expectedly, the analysis of the 1 Da channels on the peptide level is more precise than the analysis of the 6.32 mDa doublets for all resolution settings (50 000, 60 000, and 120 000), Figure 4 left panel. The CV distributions for 50 000 are clearly higher than the corresponding CV distributions accessible with 1 Da spacing. Increasing the resolution to 60 000 improves the doublets’ CVs closer to the 1 Da CV distribution, whereas with a 120 000 resolution setting, the CVs obtained from the doublets are very similar to the 1 Da spacing results. Thus, there is a tradeoff between measurement precision and scan speed. The 60 000 resolution setting seems to provide acceptable CVs for the LSF-based TMTproC proteomics, whereas 120 000 resolution generates CVs nearly indistinguishable from the established 1 Da spacing. These results are exciting, as most multiplexed proteomics experiments are currently performed with 50 000 resolution settings, and the additional 20 ms transient time seems negligible.^7,34^ We, therefore, expect SR experiments with higher plexing capability to have similar speed and sensitivity as the 1 Da TMTproC analysis.^7,18,35^ Typically, the CVs further improve when the quantitative information from peptides is integrated into the protein level. Indeed, this is also the case for our standard, where the median protein CVs improve (Figure 4, panel B).

**Figure 4.**
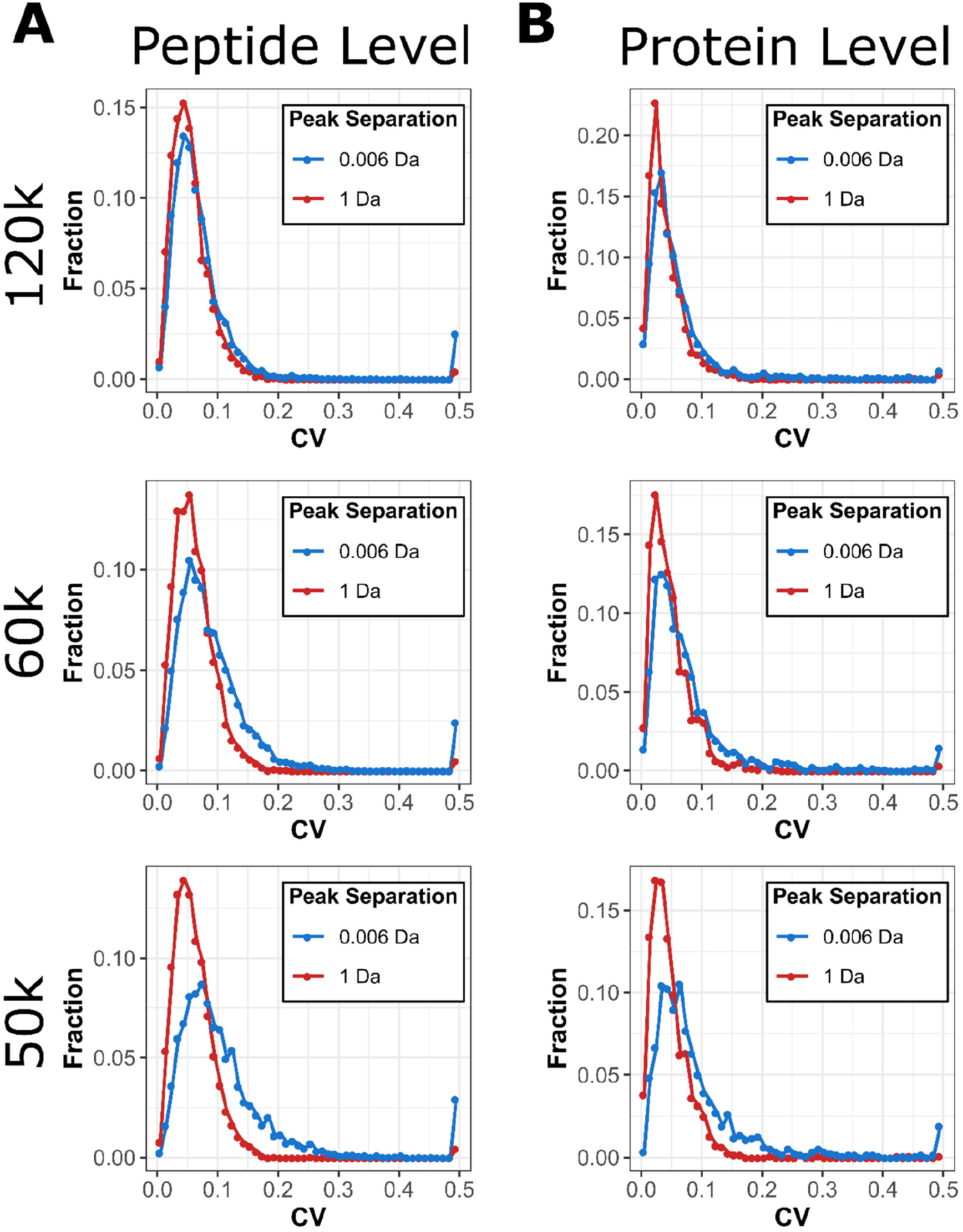
Super-resolution data processing and CVs. A) The density distributions for the CV values of quantified peptides calculated for the doublets (6.32 mDa separated channels) and singlets (1 Da separated channels) obtained in the LSF analyses of the 12-plex TMTproC LC-MS experiments with MS2 scans at the resolution settings of 120 000 (A, top panel), 60 000 (A, middle panel), and 50 000 (A, bottom panel), all defined at *m/z* 200. B) Equivalent plots once peptide quantification is integrated into protein quantification.

## Discussions

The LSF processing of as short as ∼108 ms time-domain transients (scan rate of up to 9-10 Hz, resolution setting of 50 000 at *m/z* 200 on an ultra-high-field Orbitrap instrument) resolves neutron-encoded *m/z* differences of high *m/z* complementary reporter ions in TMTc and TMTproC workflows. The time-domain transient periods are compatible with standard complement reporter ion quantification workflows, particularly because these periods allow for the parallel ion detection and accumulation capability of the modern Orbitrap instruments. Compared to conventional FT processing, this advance provides a more than 30-fold reduction in the transient period required to resolve 6.32 mDa doublets in the TMTc/TMTproC clusters. This result is in line with the specified mass difference in the doublets and the uncertainty principle definition for the SR methods in general and LSF in particular.

We have demonstrated that the time-domain transient periods increase to 256 ms (resolution setting of 120 000 at *m/z* 200) provides CV distributions for doublets (6.32 mDa channel separation) virtually indistinguishable from the 1 Da distributions. That is particularly useful for analyzing complex proteomes where high sensitivity (long ion accumulation times) is needed, as may occur in phospho-, glyco-, and single-cell proteomics.^36^ However, further increasing the time-domain transient period, for example, by 2-fold via accepting the next resolution setting of 240 000 at *m/z* 200 (512 ms periods), would reduce the efficiency of following the analyte elution and, to our understanding, is to be omitted for proteome profiling (unless ion accumulation times allow for that). For instance, analysis of the experimental data of this work demonstrates a ∼35% reduction in the number of identified peptides when time-domain transient length is increased from 256 ms to 512 ms (data not shown).

Furthermore, as frequency shifts may appear in the longer time-domain transients, keeping the ion detection times shorter is favorable. Nevertheless, these comparatively long transient times might be compatible with targeted multiplexed proteomics approaches.^37,38^ As a way forward, we envision the introduction of a TMTproC-specific intermediate resolution setting, perhaps 80 000 at *m/z* 200 (about 155 ms time-domain transient for Fusion and Q Exactive HF instruments), that could provide sufficient quantitative precision at the acceptable scan speed. We also estimate that an additional 10%-20% time-domain transient period increase may be essential to tackle the real-life proteomic samples that present a wider range of the TMTc/TMTproC channel intensities and added matrix effects.

Several experimental factors may limit the LSF (and FT) performance of the TMTc/TMTproC approach. Coulombic interaction of the ions of interest is one example. Indeed, ion-ion interaction may result in peak interference, leading to frequency shifts and, eventually, peak coalescence.^39^ These interactions can be particularly strong when ions are separated only by several mDa.^40^ Therefore, a certain care should be taken to avoid the coalescence, especially for the doublets in TMTc/TMTproC clusters, by restricting the number of elementary charges for precursor ion isolation, via the automatic gain control (AGC) setting.^41^ On the other hand, it has been shown that, thanks to the precursor ion signal split between many product ions, the coalescence may be less pronounced in a regular bottom-up proteomics experiment.^42^ Further distribution of the total charge, due to multiplexing in TMTc/TMTproC, may help to loosen the coalescence threshold. Additionally, the risks of the detrimental effects in question could increase or reduce depending on the manufacturing quality and factory tuning of the mass analyzer that is installed in a particular Orbitrap instrument.

Another limitation of the LSF/FT performance for TMTc/TMTproC, as well as other similar multiplexing strategies, is the low signal-to-noise ratios of the peaks of interest. Processing time-domain transients with low intensity ion signals may result in CVs too high for accurate protein quantitation. A related example is a TMT/TMTpro-labeled sample with a very broad range of concentrations over the (distinguishable) TMTc/TMTproC channels, so that quantitation results for the low abundance channels can be unacceptable.

Finally, high sample complexity resulting in interfering peaks in the TMTc/TMTproC cluster can be a potential limitation for the presented LSF approach when the interfering peaks are located close to the TMTc/TMTproC channels (presumably, nearby the doublets and at a distance that is less than the doublet split) or when the interfering peaks are of very large amplitude (presumably, many-fold) compared to the TMTc/TMTproC ions.

## Conclusions

Resolving the 6.32 mDa doublets in the TMTc/TMTproC clusters across the entire mass range offers a valuable increase in the multiplexing capacity of the complementary ion approach, currently from commercially available 9 to 13 channels. In addition, by modifying the distribution of heavy isotopes, it would be possible to encode 21 channels without changing the TMTpro structure.^18^ The described application of LSF in TMTc/TMTproC quantitative proteomics is a welcome addition to the arsenal of LSF-enabled strategies, such as targeted drug monitoring in mass spectrometry imaging.^43^ The described approach is compatible with other Orbitrap instruments as well, e.g. Q Exactive HF and Exploris. The LSF algorithm applied here and other SR algorithms, such as the filter-diagonalization method,^44,45^ phase-constrained spectral decomposition method,^46^ and compressed sensing, are starting to confirm the initially high expectations toward the SRMS. The described approach appears to be a sensible use of the SRMS: the doublets of interest are split by a known mass value and it is imperative that the measurement scans (transient lengths) are kept short enough to allow efficient analysis of complex proteomic mixtures.

## Acknowledgments

We are grateful for financial support through the European Research Council under grant agreement #280271 (SRMS4HESUS) and the European Horizon 2020 research and innovation program under grant agreement #964553 (ARIADNE). This work was supported by NIH grant R35GM128813, Princeton Catalysis Initiative, Eric and Wendy Schmidt Transformative Technology Fund, the U.S. Department of Energy, Office of Science, and Office of Biological and Environmental Research under award number DE-SC0018260.

## Supporting Information

Experimental and method details, including a diagrammatic representation of the experimental setup and an experimental sequence of the Fusion Lumos Orbitrap FTMS; representation of the TMTc and TMTproC basis functions for the LSF method; evaluation results of a model 4-plex TMTc workflow using UHR FTMS; examples of 6.32 mDa doublets resolution using UHR FTMS; a consistency test of 3 second aFT results versus the reference data; and evaluation results of a model 4-plex TMTc workflow: LSF vs. aFT. (pdf)

