## Supplementary Information for "Super-resolution mass spectrometry enables rapid, accurate, and highly-multiplexed proteomics at the MS2-level"

| **Figure** | **Caption** | **Page** |
| --- | --- | --- |
| S1 | Diagrammatic representation of the experimental set-up | S-2 |
| S2 | Fusion Lumos Orbitrap FTMS experimental sequence | S-3 |
| S3 | TMTproC and TMTc basis functions for the LSF method | S-4 |
| S4 | Evaluation of a model 4-plex TMTc workflow using UHR FTMS | S-5 |
| S5 | Examples of 6.32 mDa doublets resolution using UHR FTMS | S-6 |
| S6 | Consistency test of 3-s aFT results versus the reference data | S-7 |
| S7 | Evaluation of a model 4 -plex TMTc workflow: LSF vs. aFT | S-8 |

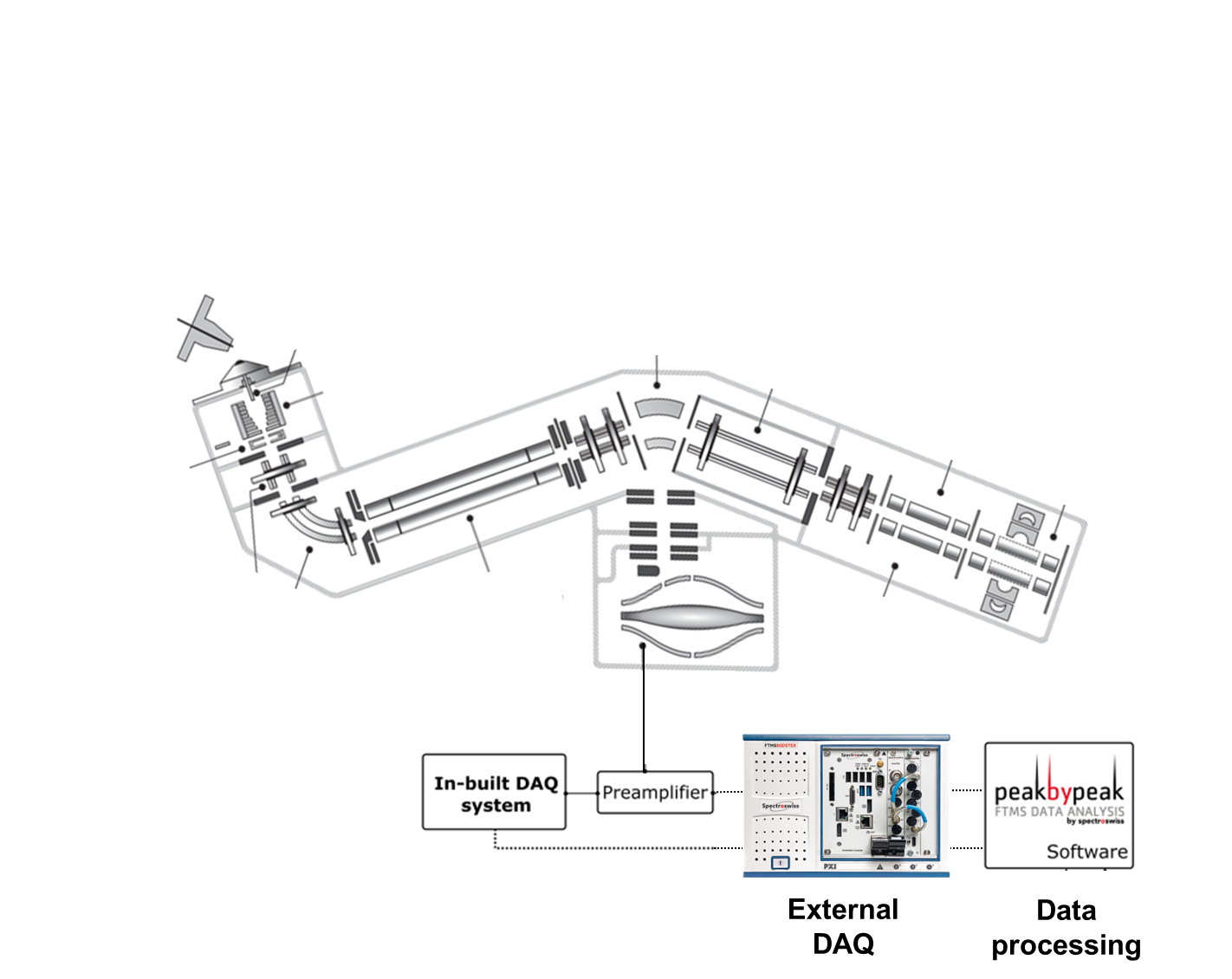

**Figure S1.** The TMTc/TMTproC quantitative proteomics workflow employed in the current work. Diagrammatic representation of the components of the Fusion^TM^ Lumos^TM^ Orbitrap^TM^ FTMS coupling along with the external DAQ system (FTMS Booster X2) and the allied data processing software (on the basis of Peak-by-Peak software suite).

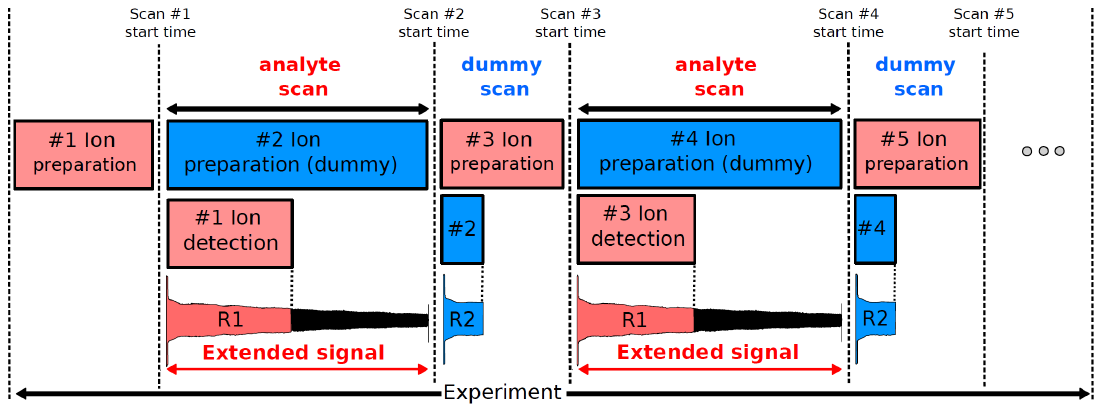

**Figure S2.** Fusion^TM^ Lumos^TM^ Orbitrap^TM^ FTMS experimental sequence that was used to generate the datasets with the extended transient periods. The sequence takes advantage of the parallel ion detection/accumulation capability of the modern Orbitrap instruments, in order to generate extended-length transients. Therefore, enabling the extended period transients can be achieved with method optimization and does not require hardware modification.

The introduction of the dummy scans extends the duration of the “extended signal” for the previous scan. More specifically, the final period of the extended time-domain signal will depend on the ion preparation event duration of the dummy scan. Users can control the ion preparation duration by manipulating with ion accumulation and/or ion fragmentation events in the dummy scans.

The built-in DAQ system of an Orbitrap will record time-domain transient from a Start trigger signal and for a period specified prior to the measurements by setting the resolution target, e.g., 256 ms for resolution setting of 120 000 at *m/z* 200. To record the full, extended time-domain transient, the external high-performance DAQ system is required to record time-domain signals from a Start trigger (ion injection into the Orbitrap mass analyzer) to the Stop trigger (ion ejection from the Orbitrap mass analyzer). The period of the extended signal can be arbitrary – there is no requirement to always record time-domain transients with factor of 2 increase in length.

At the end of the measurements, the dummy scans are excluded from the consideration and only the analyte scans with extended signals are processed.

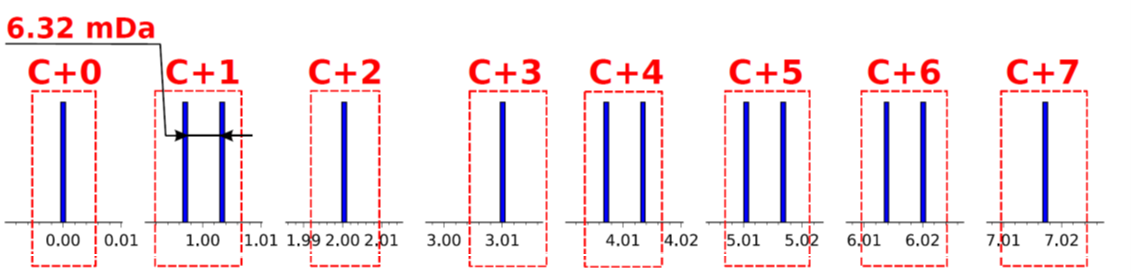

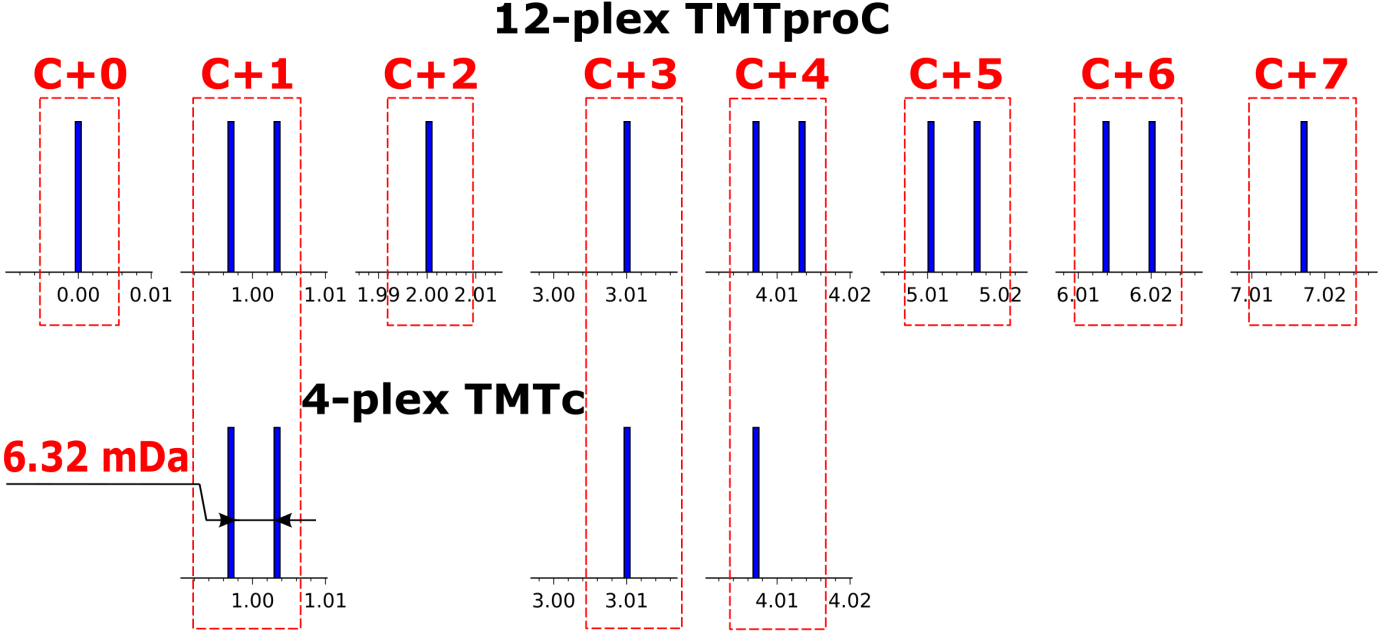

**Figure S3.** Basis functions for the LSF method, corresponding to 12 TMTproC channels with four 6.32 mDa doublets and four singlets (top panel), and 4 TMTc channels with two singlets and one 6.32 mDa doublet (bottom panel).

.

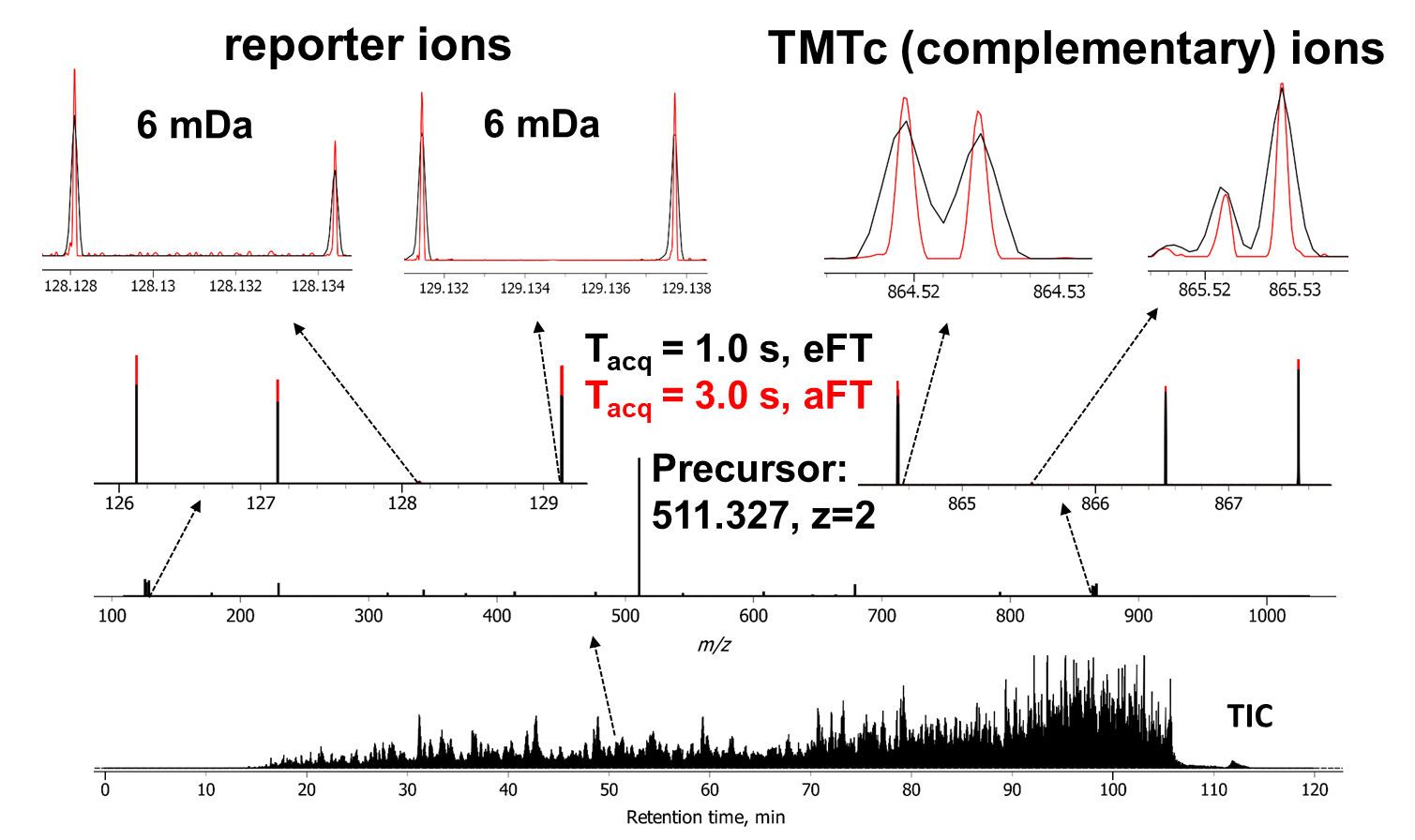

**Figure S4.** Evaluation of a model 4-plex TMTc workflow using UHR Orbitrap FTMS. The LC-MS/MS quantitative proteomics experiments were performed with a Fusion Lumos Orbitrap FTMS equipped with an external high-performance data acquisition system, FTMS Booster X2 (Spectroswiss). The MS/MS experiments were performed using HCD.

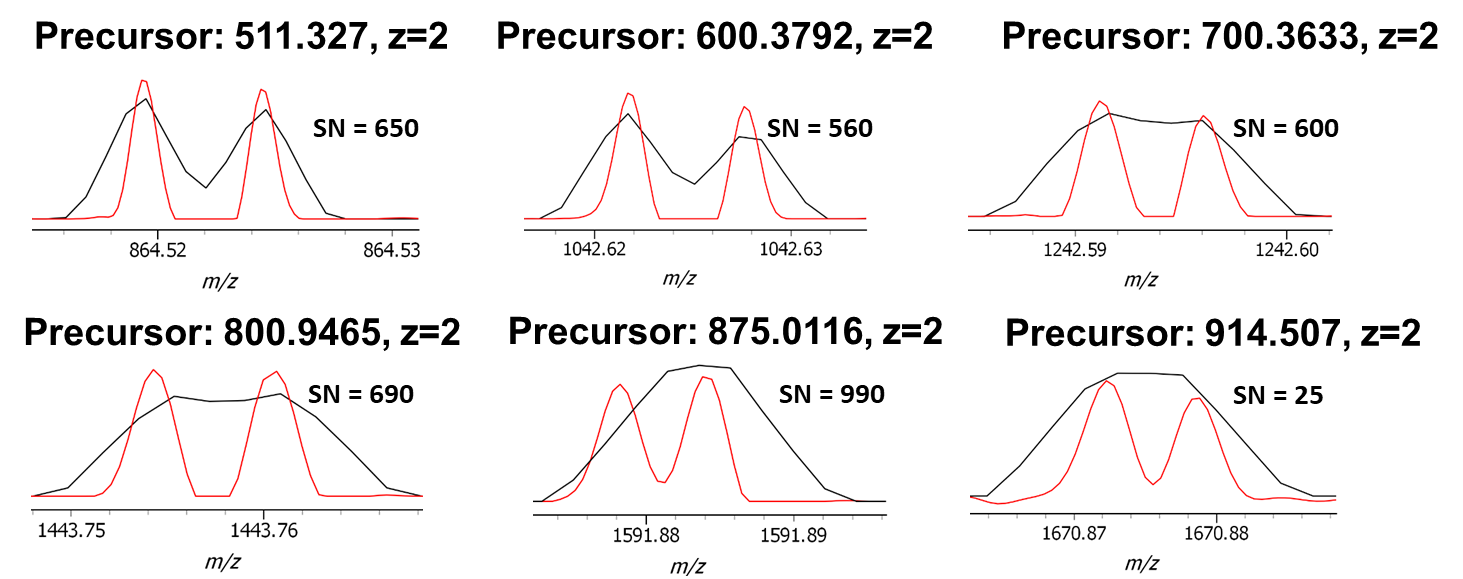

**Figure S5.** Examples of 6.32 mDa doublets resolution upon evaluation of a model 4-plex TMTc workflow using UHR Orbitrap FTMS. Experiments were performed with a Fusion Lumos Orbitrap FTMS equipped with an external high-performance data acquisition system, FTMS Booster X2 (Spectroswiss). The MS/MS experiments were performed using HCD. Compared are the mass spectra obtained with the maximum supported resolution capability of the FTMS instrument (resolution setting 500 000 at *m/z* 200, transient period 1 s, eFT processing, data shown in black) and the UHR results (FTMS Booster X2, transient period 3 s, aFT processing, data shown in red).

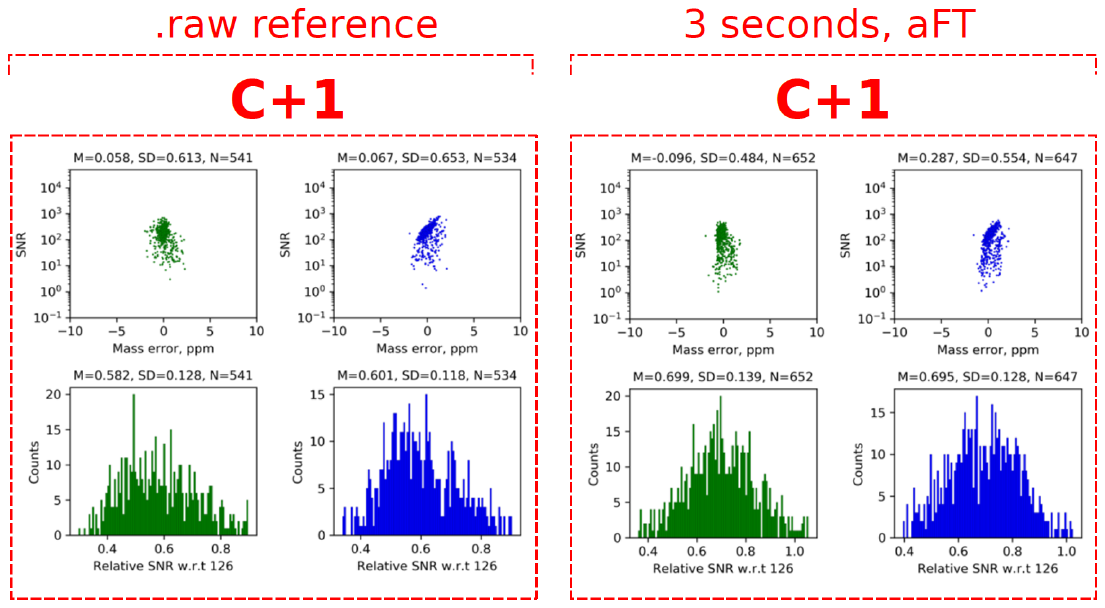

**Figure S6.** Consistency test of the 3-seconds aFT results versus the .raw reference data, using the ions from the C+1 doublet in the analysis of the 4-plex TMTc data. The S/N vs. mass error scatter plots (top panels) and the corresponding density distributions for the abundances relative to the abundance of the C+4 singlet (bottom plots) are shown for data with S/N>1 and mass error range of ±10 ppm.

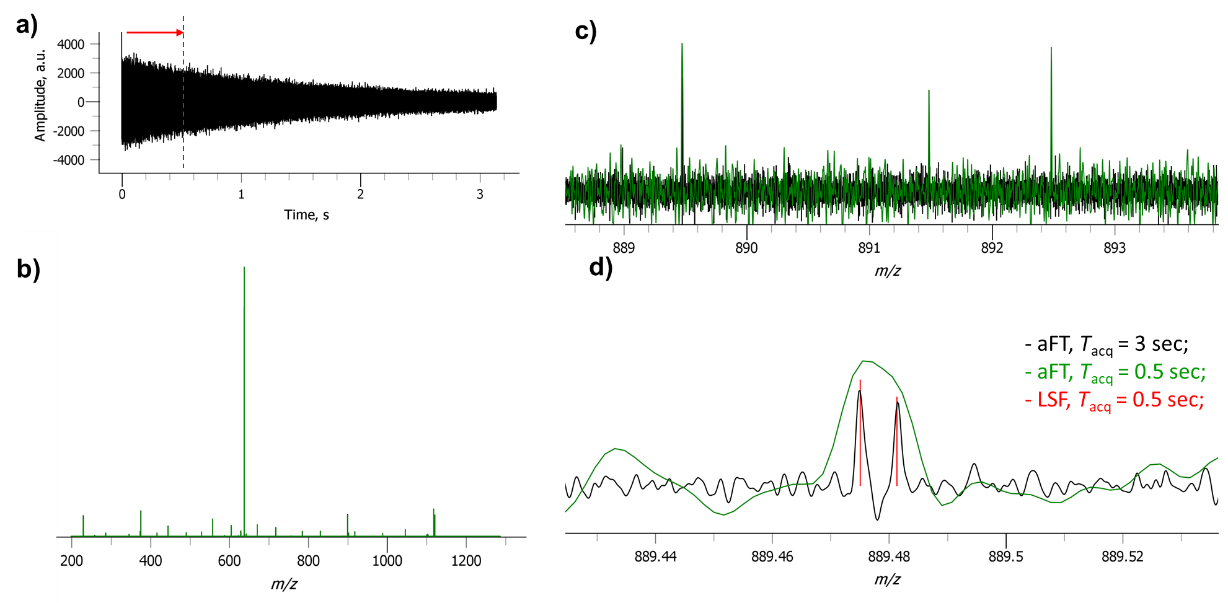

**Figure S7.** Performance evaluation of a model 4 -plex TMTc workflow using UHR Orbitrap FTMS. Experiments were performed with a Fusion Lumos Orbitrap FTMS equipped with an external high-performance data acquisition system, FTMS Booster X2 (Spectroswiss). The MS/MS experiments were performed using HCD. The selected example shows: (a) a 3 s time-domain transient of an HCD MS/MS of a doubly charged precursor ion at 638.386 *m/z* acquired with an external high-performance data acquisition system (FTMS Booster X2, Spectroswiss); (b) a corresponding tandem spectrum of a doubly charged precursor ion at 638.386 *m/z* produced by absorption mode FT (aFT); (c) an expanded view into the tandem mass spectrum showing the region of a TMTc cluster; and (d) a further expanded view into the 6.32 mDa doublet comparing results of different data processing routines: aFT of 3 s transient (shown in black), aFT of 1 s transient (shown in green), and LSF of 0.5 s transient (shown in red).
